# Functional Exploration of Heterotrimeric Kinesin-II in IFT and Ciliary Length Control in *Chlamydomonas*

**DOI:** 10.1101/2020.05.14.096529

**Authors:** Shufen Li, Wei Chen, Kirsty Y Wan, Hui Tao, Xin Liang, Junmin Pan

## Abstract

Heterotrimeric organization of kinesin-II is essential for its function in anterograde IFT in ciliogenesis. However, the molecular basis of forming this complex for its function is not well understood. In addition, the anterograde IFT velocity varies significantly in different organisms, but how motor speed affects ciliary length is not clear. We show that *Chlamydomonas* kinesin-II (CrKinesin-II) involves distinct mechanisms from mammals and C. *elegans* in its assembly to necessitate its function in IFT. Furthermore, chimeric CrKinesin-II with human kinesin-II motor domains functioned *in vitro* and *in vivo*, leading to a ~2.8-fold reduced anterograde IFT velocity and a similar fold reduction in IFT injection rate that supposedly correlates with ciliary assembly activity. However, the ciliary length was only mildly reduced (~15%). Modelling analyses suggest that such a non-linear scaling relationship between IFT velocity and ciliary length can be accounted for by limitation of the motors and/or its ciliary cargoes, e.g. tubulin.

## INTRODUCTION

It is well-established that cilia are conserved cellular organelles that play pivotal roles in signaling and cellular motilities, and defects in cilia are linked with numerous human diseases and developmental disorders (Anvarian et al., 2019; Bangs and Anderson, 2017; Reiter and Leroux, 2017). The assembly and maintenance of cilia requires intraflagellar transport (IFT), a bidirectional movement of protein complexes (IFT complexes) between the ciliary membrane and the axoneme (Kozminski et al., 1993). The anterograde transport (from ciliary base to tip) is driven by kinesin-2 whereas retrograde transport (from ciliary tip to base) is powered by cytoplasmic dynein 2/1b (Rosenbaum and Witman, 2002; Scholey, 2003). The IFT complexes, which consist of IFT-A and IFT-B complexes, serve as cargo adaptors to recruit ciliary proteins (Lechtreck, 2015; Taschner and Lorentzen, 2016), and are assembled into linear arrays termed IFT particles or IFT trains (Kozminski et al., 1993; Pigino et al., 2009).

Heterotrimeric kinesin-2 (kinesin-II) is essential for anterograde IFT and ciliogenesis in most ciliated cells while both heterotrimeric and homodimeric kinesin-2 collaboratively drive anterograde IFT in *C. elegans* (Scholey, 2013). In contrast to most kinesins with two identical motor subunits, kinesin-II consists of two non-identical motor subunits and one non-motor subunit (KAP) (Hirokawa et al., 2009; Verhey and Hammond, 2009). The heterotrimeric organization of kinesin-II is required for IFT because mutation in either subunit abolishes or impairs IFT in various organisms (Engelke et al., 2019; Kozminski et al., 1995; Liang et al., 2014; Lin et al., 2003; Miller et al., 2005; Mueller et al., 2005; Nonaka et al., 1998; Snow et al., 2004). Though different systems use similar machinery, the underlying mechanism of this heterotrimeric organization for IFT appear to diversify in the worm and mammals. For example, homodimer of kinesin-II motor subunits cannot be formed and motors with two identical motor domains are not functional *in vivo* in worm (Brunnbauer et al., 2010; Pan et al., 2010) whereas in mammalian cells, kinesin-II subunit KIF3A can form homodimers but cannot associate with IFT complexes (Funabashi et al., 2018). It is intriguing to learn how this heterotrimeric organization requirement is conserved and diversified, especially in the unicellular eukaryote *Chlamydomonas* in which IFT was first discovered (Kozminski et al., 1993).

During IFT, the velocity of anterograde IFT driven by kinesin-II varies with the types of organisms. Though the assay conditions would slightly affect the measurements, the velocity of anterograde IFT is ~2.2 μm/s in *Chlamydomonas* and *Trypanosome* (Bertiaux et al., 2018a; Brown et al., 2015; Dentler, 2005; Engel et al., 2009; Liang et al., 2014; Wingfield et al., 2017). In contrast, mammalian cells and worms have a much slower velocity (~0.5 μm/s) (Broekhuis et al., 2014; Engelke et al., 2019; Follit et al., 2006; Snow et al., 2004). Notably, *Chlamydomonas* and *Trypanosoma* have longer cilia whereas mammalian cells tend to have shorter cilia. It is intriguing how the motor speed affects IFT and ciliary assembly. In addition, kinesin-II is thought to drive IFT complexes from the ciliary base through the transition zone to the cilium (Prevo et al., 2015; Scholey, 2013; Wingfield et al., 2017). However, it was also argued that kinesin-II takes over anterograde transport only after IFT complexes enter cilia via a mechanism where the mechanochemical coupling of kinesin-2 is dispensable (Nachury and Mick, 2019). Thus, it is interesting to know how kinesin-II motor activity affects entry of IFT trains into cilia.

In this work, we reveal distinct mechanisms for the requirement of the heterotrimeric organization of *Chlamydomonas* kinesin-II (CrKinesin-II), demonstrating that a similar machinery may employ divergent mechanisms for the function. Furthermore, we generated chimeric CrKinesin-II with motor domains of human kinesin-II (HsKinesin-II) and show that it can perform motility function *in vitro* and *in vivo* but with an ~2.8-fold reduction in the velocity of motor and/or anterograde IFT. The reduced motor velocity results in a similar reduction in the IFT injection rate, suggesting that kinesin-II activity is involved in moving IFT complexes into cilia. IFT injection rate has been shown to correlate with ciliary assembly activity and thus ciliary length (marshall 2001, 2005 2009). However, the effect of changing motor speed on ciliary length has not been directly demonstrated *in vivo* until now. Interestingly, our results reveal that the ciliary length of the cells expressing the slow chimeric motor is only mildly reduced (~15%). Using a modeling approach to understand the effect of motor speed on ciliary assembly and length, we reveal that limitation of motors or key ciliary components are likely the key determinants of ciliary length.

## RESULTS

### The requirement of heterotrimeric organization of CrKinesin-II for IFT

The function of kinesin-II in IFT requires two non-identical motor subunits and a non-motor subunit, kinesin-associated protein (KAP). To understand how this organization is required for the function of CrKinesin-II in IFT, we analyzed whether CrKinesin-II with two identical motor domains can coordinate for motility *in vitro* and *in vivo*, and whether each of the motor subunits can interact with KAP independently without the other subunit. For *in vitro* motility assay, we generated CrKinesin-II constructs with fluorescent tags as well as tags for protein purification as indicated (Figure 1A). To generate CrKinesin-II constructs with two identical motor domains, the motor domain of FLA8 was replaced with that of FLA10 and *vice versa* (Figure 1B). The recombinant wild-type CrKinesin-II as well as the chimeric motors FLA10/FLA10’/KAP and FLA8/FLA8’/KAP were expressed respectively in SF9 cells and purified (Figure S1).

**Figure 1.**
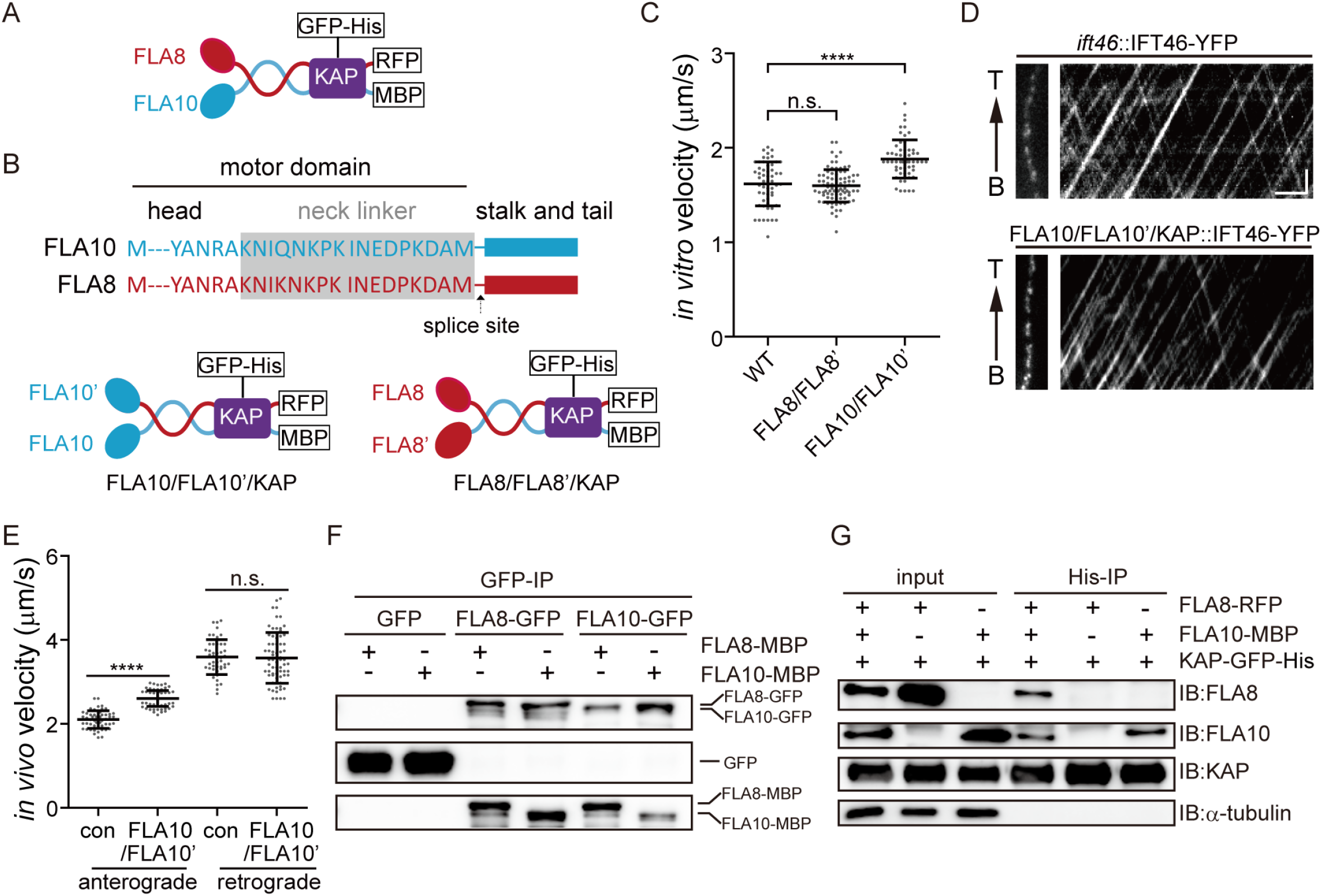
Requirement of the heterotrimeric organization of CrKinesin-II for IFT (See also Figure S1-3). (A) Schematic diagram of recombinant CrKinesin-II for expression/purification. (B) Overview of chimeric CrKinesin-II constructs with two identical motor domains. The motor domain of FLA10 was replaced with that of FLA8 or vice versa to create chimeric kinesin-IIs with two identical motor domains. Arrow indicates the splice site after the neck linker (gray) for creating the chimeric constructs. (C) *In vitro* motility assay of chimeric CrKinesin-IIs with two identical motor heads at 23 °C. Please note, KAP is present in the chimeric motors. Data shown are mean ± SD. ****p < 0.0001; n.s., statistically not significant. (D-E) Analysis of IFT. The velocities of IFT46-YFP expressed in FLA10/FLA10’/KAP cells or in an *ift46* rescue strain expressing IFT46-YFP (as a control) were measured using TIRF microscopy. Representative kymographs of IFT (D) and the measurements (E). Data shown are mean ± SD. ****p < 0.0001; n.s., statistically not significant. (F) Self-interaction of FLA10 or FLA8. FLA10-GFP and FLA10-MBP or FLA8-GFP and FLA8-MBP were co-expressed respectively in 293T cells followed by immunoprecipitation with anti-GFP antibody and immunoblotting with GFP and MBP antibodies, respectively. (G) FLA10 interacts with KAP while FLA8 does not. FLA10-MBP or FLA8-RFP was co-expressed respectively with KAP-GFP-His followed by pull-down with a Ni column and immunoblotting with the indicated antibodies.

We used total internal reflection fluorescence (TIRF) microscopy to determine the motility of the purified motors. Compared to wild type motors (1.62 ± 0.23 μm/s, n = 55), FLA8/FLA8’/KAP (1.60 μm/s ± 0.17, n = 83) moved with a similar velocity while FLA10/FLA10’/KAP (1.88 ± 0.20 μm/s, n = 56) showed a slightly higher velocity. Thus, chimeric motors with two identical motor domains (i.e. FLA10/FLA10 and FLA8/FLA8) of Crkinesin-II can functionally coordinate. These results were consistent with the reports for kinesin-II from *C. elegans* and mammal (Brunnbauer et al., 2010; Muthukrishnan et al., 2009; Pan et al., 2010). However, kinesin-II with two identical motor domains of KLP20 in *C. elegans* did not function *in vivo* (Pan et al., 2010). It was intriguing whether this was conserved in *Chlamydomonas*. Thus, we tested the *in vivo* functionality of the chimeric motor FLA10/FLA10’/KAP as FLA10 and KLP20 are homologous. To this end, *FLA10’-HA* was transformed into an aflagellate *fla8* mutant and the transformants were expected to form a chimera with two FLA10 motor domains *in vivo*. FLA10/FLA10’/KAP transformants rescued the aflagellar phenotype of *fla8* in terms of ciliary length and ciliary regeneration kinetics (Figure S2), indicating that Crkinesin-II with two identical motor domains of FLA10 performs proper physiological function *in vivo*. To determine whether the transformants indeed rescued IFT, IFT46-YFP was expressed respectively in FLA10/FLA10’/KAP cell and an *ift46* mutant (as a control) (Lv et al., 2017). The retrograde IFT in FLA10/FLA10’/KAP cells (3.57 ± 0.60 μm/s, n = 70) showed a similar velocity to that in the control cells (3.59 ± 0.42 μm/s, n = 46). The velocity of anterograde IFT in FLA10/FLA10’ (2.60 ± 0.19 μm/s, n = 73) was slightly higher relative to the control (2.10 ± 0.21 μm/s, n = 53) (Figure 1D and E). Thus, CrKinesin-II with two identical motor domains of FLA10 could function *in vivo*, which is in contrast to the results in *C. elegans* (Pan et al., 2010).

We next examined whether FLA10 or FLA8 was able to form homodimer and interact with KAP. FLA10-GFP and FLA10-MBP were co-expressed in HEK293T cells followed by immunoprecipitation with an anti-GFP antibody and immunoblotting with GFP and MBP antibodies, respectively. Similar experiments were performed for FLA8. Both FLA10 and FLA8 could self-interact (Figure 1F). Supposing that self-interaction of FLA10 or FLA8 can form proper homodimer, we then asked whether they could interact with KAP. FLA10-MBP and FLA8-RFP were co-expressed with KAP-GFP-His, respectively, followed by pull-down with Ni beads and immunoblotting (Figure G). Interestingly, FLA10-MBP interacted with KAP while FLA8-RFP did not. KAP is required for kinesin-II’s full activation and recruitment to ciliary base (Mueller et al., 2005; Sonar et al., 2020) and FLA8 homologue KIF3B is required for the interaction of kinesin-II with IFT complex (Funabashi et al., 2018). Thus, neither homodimers of FLA10 or FLA8 can function in IFT, because it is likely that the FLA10 homodimer could not interact with IFT complex while FLA8 homodimer could not interact with KAP, which explains the necessity of a heterotrimetric organization of CrKinesin-II for IFT. We showed that both FLA10 and FLA8 could likely form homodimers *in vitro*, however, formation of homodimers *in vivo* is expected to interfere with the proper formation of heterotrimeric kinesin-II. We found that FLA10 in *fla8* mutant was undetectable likely due to protein degradation (Figure S3), indicating that homodimer formation could not occur *in vivo*.

### Chimeric CrKinesin-II with motor domains of HsKinesin-II functions *in vitro* and performs physiological function *in vivo* of *Chlamydomonas*

Kinesin-II functions in various ciliated organisms to drive anterograde IFT. However, it has quite different properties. For example, the motility of kinesin-II and thus that of the anterograde IFT vary several folds between *Chlamydomonas* and mammal (Broekhuis et al., 2014; Brown et al., 2015; Engelke et al., 2019; Follit et al., 2006; Kozminski et al., 1993; Muthukrishnan et al., 2009; Wingfield et al., 2017). We wanted to examine whether a chimeric kinesin-II motor with motor domains from different species could perform motility function and what would be the physiological consequences. To this end, we generated chimeric CrKinesin-IIs with one or two motor domains of human kinesin-II (Figure S4). Wild type and chimeric kinesin-IIs were expressed respectively in SF9 cells and subsequently purified (Figure 2 and Figure S4). *In vitro* motility assay showed that all the chimeras indeed could move. However, they had a similar motility to that of HsKinesin-II and was significantly slower than CrKinesin-II (~3-fold reduction) (Figure 2B). This result suggests that motor domains from different species can coordinate and the slower motor subunit determines the velocity of the chimeric motor.

**Figure 2.**
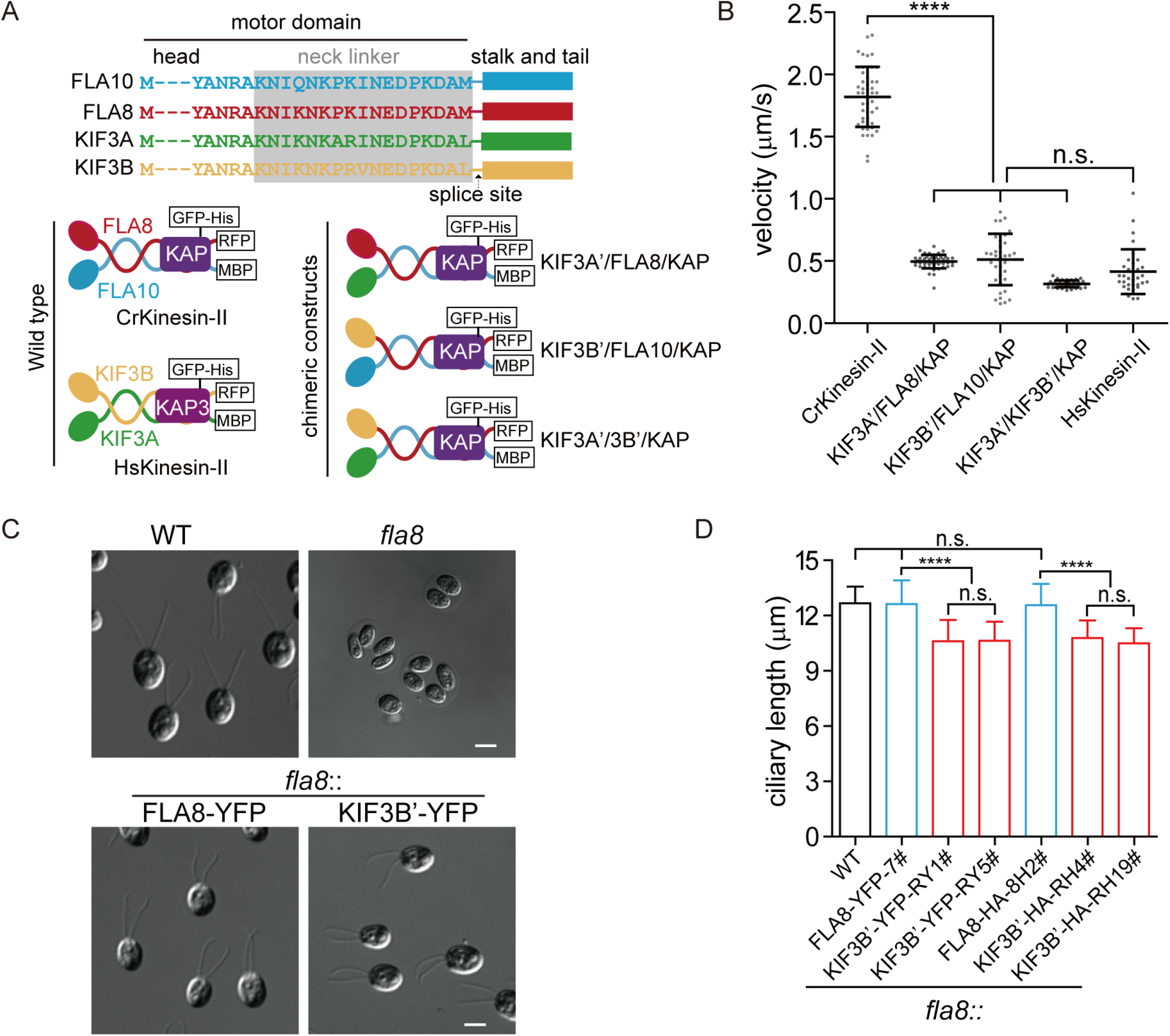
Chimeric CrKinesin-IIs with motor domains of HsKinesin-II function *in vitro* and *in vivo* (see also Figure S4). (A) Overview of chimeric CrKinesin-II constructs. The motor domains of FLA10, FLA8 or both in CrKinesin-II were replaced with their counterparts of HsKinesin-II, respectively. (B) *In vitro* motility assay of CrKinesin-II chimeras at 23 °C. The rates are the following: 1.82 ± 0.24 μm/s (n = 48) for CrKinesin-II; 0.50 ± 0.05 μm/s (n = 50) for KIF3A’/FLA8/KAP; 0.51 ± 0.20 μm/s (n = 37) for KIF3B’/FLA10/KAP; 0.32 ± 0.03 μm/s (n = 40) for KIF3A’/KIF3B’/KAP and 0.41 ± 0.18 μm/s (n = 48) for HsKinesin-II. ****p < 0.0001; n.s., statistically not significant. (C) Rescue of the aflagellate phenotype of *fla8* by *FLA8-YFP* or *KIF3B’-YFP*. *fla8* was transformed with *FLA8-YFP* and *KIF3B’-YFP* respectively. Cells were imaged using differential interference contrast microscopy. Wild type (WT) and *fla8* cells were shown as control. Bar, 5 μm. (D) Cells expressing KIF3B’/FLA10/KAP chimera show robust but mild decrease in ciliary length. The ciliary length in steady state cells as indicated were measured.

Though the chimeric motors could function *in vitro*, it remains a question whether it can fulfill its physiological functions *in vivo*. Furthermore, the ~3-fold slower speed of the chimeric Crkinesin-II compared to the wild type CrKinesin-II would also allow us to examine how the velocity of the motor contributes to ciliary length and regeneration. We chose to test the chimeric CrKinesin-II KIF3B’/FLA10/KAP in *Chlamydomonas*. To do this, the *fla8* mutant was transformed with *KIF3B’-YFP* or *FLA8-YFP* (as a control). The transformants were expected to form KIF3B’-YFP/FLA10/KAP or FLA8-YFP/FLA10/KAP motors. Examination of the ciliary phenotype revealed that both transformants rescued the aflagellar phenotypes of *fla8* (Figure 2C), indicating that the chimeric KIF3B’-YFP/FLA10/KAP could function *in vivo*. Next, we measured ciliary length. The cilia in the KIF3B’-YFP/FLA10/KAP cells had an average length of 10.6 ± 1.1 μm (n=50), ~15% shorter compared to the control cells (12.6 ± 1.3 μm, n=50) and wild type cells (Figure 2D). We further verified this change by using *fla8* cells expressing *KIF3B’-HA*, which again showed ~15% reduction in length, and *FLA8-HA*, which rescues the ciliary length to the control level (Figure 2D). These observations demonstrate that although the reduction in ciliary length was mild, it is a robust consequence of slow IFT mediated by a slower kinesin-II motor. Taken together, we showed that chimeric CrKinesin-II with motor domain of HsKinesin-II could function *in vitro* and *in vivo* though the chimeric motor did not fully recover the ciliary phenotype.

### Chimeric CrKinesin-II with human motor domain results in a significant reduction in IFT injection rate

The recovery of ciliary phenotype in KIF3B’-YFP/FLA10/KAP cells suggests that the chimeric motor KIF3B’-YFP/FLA10/KAP functions in anterograde IFT. Because this chimeric motor is slower *in vitro* than the wild type motor, we first examined the motor velocity *in vivo*. Though the velocities of KIF3B’-YFP (~0.91 μm/s) and FLA8-YFP (~2.61 μm/s) were higher than their *in vitro* data respectively (Figure 3A and Figure 2B), KIF3B’-YFP was ~2.8-fold slower relative to FLA8-YFP, which is consistent with the *in vitro* assay data (Figure 2B). The motor speed should reflect the velocity of anterograde IFT. This was confirmed by measuring the velocity of an IFT protein (IFT46-YFP) in *fla8* mutants that were transformed with HA-tagged *FLA8* or *KIF3B’* (Figure 3B). The anterograde velocities of IFT46-YFP in the FLA8 transformant (2.29 ± 0.25 μm/s, n = 61) and KIF3B’ transformant (0.80 ± 0.08 μm/s, n = 61) were similar to the velocities of the motors. Based on these results, we conclude that the chimeric CrKinesin-II KIF3B’ /FLA10/KAP function in IFT but with a slower speed.

**Figure 3.**
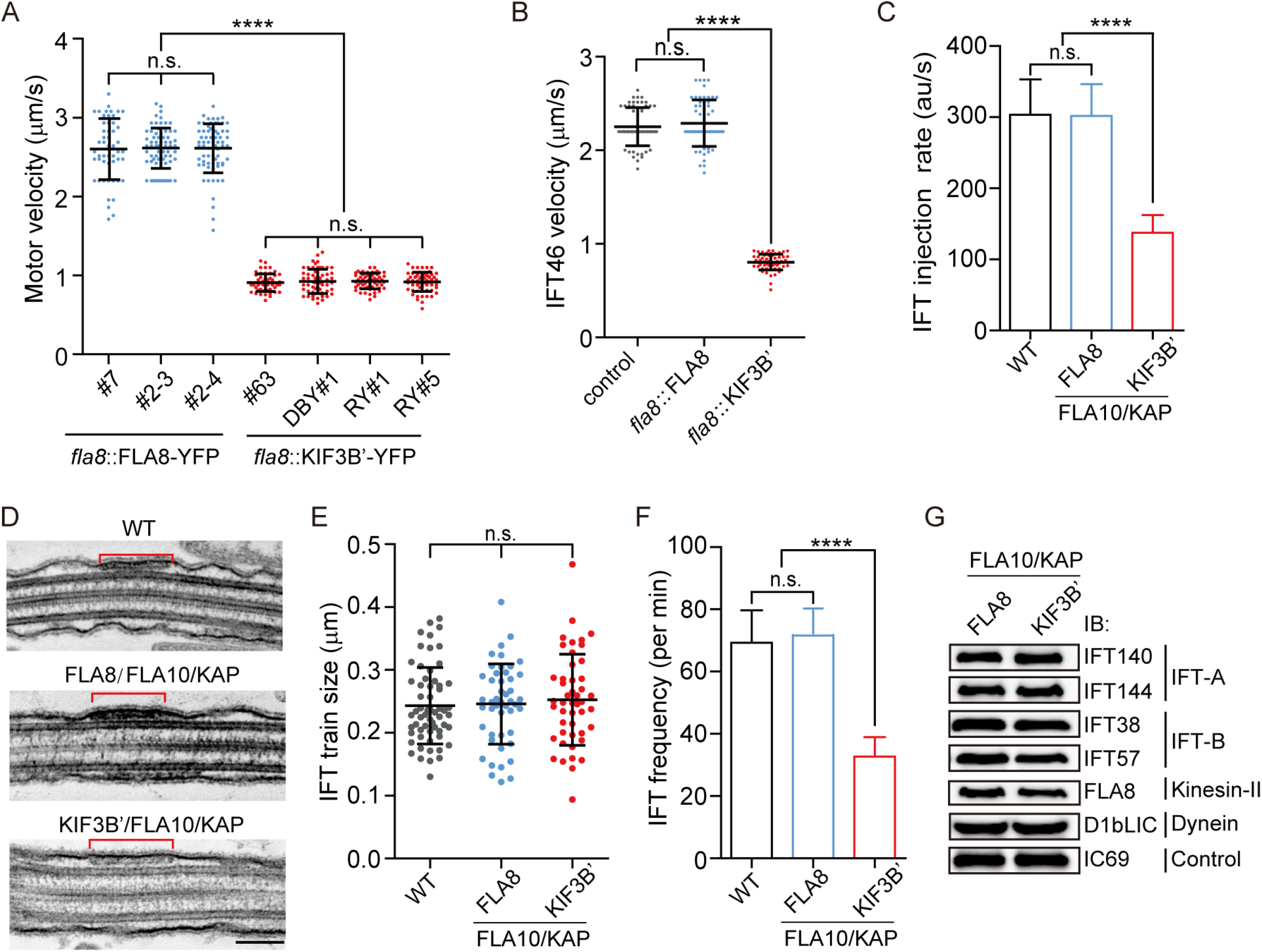
Chimeric KIF3B’/FLA10 motor leads to significant reduction in IFT injection rate but slight decrease in ciliary length. (A) Velocities of FLA8-YFP and KIF3B’-YFP. The anterograde velocities of FLA8-YFP and KIF3B’-YFP that were expressed respectively in *fla8* cells were assayed using TIRF microscopy. ****p < 0.0001; n.s., statistically not significant. (B) Anterograde velocities of IFT46-YFP in FLA8 and KIF3B’ transgenic cells. *IFT46-YFP* was transformed into *fla8* cells expressing FLA8-HA or KIF3B’-HA followed by analysis using TIRF microscopy. *ift46* cells expressing IFT46-YFP were used as a control. (C) KIF3B’/FLA10/KAP chimera leads to ~2.2-fold reduction in IFT injection rate, which was measured by monitoring fluorescence intensity of IFT46-YFP that enters into cilia per time using TIRF microscopy. (D-E) Analysis of the IFT train size. Representative TEM images of cilia showing IFT trains (D) and the average train size was similar among the indicated samples (E). Bar, 100 nm. (F) KIF3B’/FLA10 chimera leads to ~2.1-fold reduction in IFT frequency. *IFT46-YFP* was expressed in the indicated cells followed by analysis using TIRF microscopy. 69.3 ± 10.34 min^−1^ (n = 60) for wild type (WT), 72 ± 8.2 min^−1^ (n = 60) for FLA8-HA/FLA10/KAP and 32.7 ± 6.24 min^−1^ (n = 60) for KIF3B’-HA/FLA10/KAP cells. (G) Cells expressing chimeric KIF3B’/FLA10/KAP have similar ciliary levels of IFT proteins to the controls. The cilia were isolated from the indicated cells. Equal amounts of ciliary proteins were analyzed by immunoblotting with the indicated antibodies.

We next analyzed how motor activity change influences the ciliary entry of IFT trains into the cilium. The IFT injection rates, the amount of IFT trains entering cilia per unit time (au/s), from cells expressing chimeric and wild type kinesin-II were measured. Using TIRF microscopy, we estimated IFT injection rate by monitoring the amount of IFT46-YFP entered into cilia per unit time in KIF3B’-HA/FLA10/KAP cells; FLA8-HA/FLA10/KAP and wild type cells were as control. The IFT injection rate in FLA8-HA/FLA10/KAP (301.78 ± 44.21 au/s, n = 60) was similar to that in the wild type cells (303.35 ± 49.42 au/s, n = 60) and was 2.2-fold of that in KIF3B’-HA/FLA10/KAP (137.57.35 ± 24.62 au/s, n = 60) (Figure 3C), a similar fold-change as the IFT velocity.

We then wondered how the IFT injection rate was reduced. Intuitively, IFT injection rate is the product of IFT injection frequency (number of IFT trains entering cilia per unit time) and the average size of IFT trains. Using transmission electron microscopy, we found that the average train size was similar among FLA8/FLA10/KAP, KIF3B’/FLA10/KAP and wild type cells (243 – 253 nm) (Figure 3D-E), which is consistent with a previous report (Stepanek and Pigino, 2016). In contrast, the IFT frequency of KIF3B’/FLA10/KAP cells was reduced by ~2.1-fold as measured by TIRF microscopy (Figure 3F), which is similar to the fold-reduction in IFT injection rate (Figure 3C). Thus, we conclude that the reduction in IFT frequency accounts for the reduction in IFT injection rate in KIF3B’-HA/FLA10/KAP cells.

It is intriguing how the amount of IFT proteins inside the cilium is changed given the change in IFT injection rate and IFT velocity in the cells expressing chimeric kinesin-II. The amount of IFT protein in a cilium is given by the following equation if retrograde IFT is not considered: *M* = *L*/*v* × *J*, where *M* is the quantity of IFT proteins in a cilium; *L* is the ciliary length (μm); *v* is the velocity of anterograde IFT (μm/s) and *J* is IFT injection rate (s^−1^). Compared to the control cells, the ciliary length in the chimeric motor cells is about 15% shorter, the IFT injection rate and velocity were reduced ~2.8 and ~2.2-fold respectively. Given these compensatory contributions, we predict that the ciliary levels of IFT proteins should be similar between these two cases. We performed immunoblotting with isolated cilia to confirm that the ciliary levels of IFT proteins were indeed similar (Figure 3G), supporting the above mentioned reasoning. Taken together, we showed that the chimeric kinesin is functional in IFT though with a reduced velocity and it significantly reduces the IFT injection rate by down-regulating IFT injection frequency.

### Modeling: relationship between motor speed, ciliary assembly and length control

The eukaryotic cilium is a model system for probing the phenomenon of organellar size control and equilibration (Chan and Marshall, 2012). Several models have been proposed to explain ciliary length control, specifically in the *Chlamydomonas* system (Bertiaux et al., 2018b; Fai et al., 2019; Hendel et al., 2018; Ludington et al., 2015; Marshall and Rosenbaum, 2001; Patra et al., 2020; Wemmer et al., 2020). Our results show that a motor with ~2.8-fold reduction in speed results in a small change in ciliary length (~15% shorter). To understand the relationship between the IFT velocity and ciliary length, we turned to a modelling approach. We first considered a well-established phenomenological model for ciliary length control (Marshall and Rosenbaum, 2001). In this simplest case, it is assumed that IFT limits cilia regeneration, leading to an empirical inverse scaling law between IFT injection rate and cilium length (Engel et al., 2009). The reduction in IFT injection rate during ciliary elongation results in decreased ciliary assembly activity, which is eventually balanced with a constant disassembly rate leading to a final steady-state length. However, this model predicts a linear dependence between the steady-state ciliary length and IFT velocity. This is inconsistent with our data (Figure 2D), suggesting that more detailed aspects of the IFT dynamics must be incorporated to explain our observation.

To this end, we turned to more recently published models and extended the analysis to explore the dependence of ciliogenesis dynamics in different physical regimes to understand how motor speed affects ciliary length (see Methods). These belong to a class of ‘diffusion’ models (Fai et al., 2019; Hendel et al., 2018) in which there are two key assumptions. First, that over ciliary regeneration timescales, there is a limiting pool of resources, in this case of kinesin motors (and hence also of free tubulin). Second, diffusion limits the rate at which kinesin motors can return back from the ciliary tip to the base, thereby resulting in limitation of motors (Chien et al., 2017). We assume that kinesin-II motors transport anterograde IFT trains ballistically with speed *v*, deposit cargoes including tubulin subunits at the growing ciliary tip, and then diffuse steadily back to the base. Using this interpretation, we now obtain a highly nonlinear dependence of the fold-change in speed *v*/*v*_0_ versus the fold-change in cilium length *L*/*L*_0_, where *v*_0_, *L*_0_ denote the wild type motor speed and final cilium lengths respectively. For realistic parameters and a reduced motor speed of ~*v*_0_/3, the model predicts a ~18% reduction in *L* (Figure 4A), which qualitatively agrees with our data (Figure 2D). More generally, this model predicts that a faster motor speed would lead to a very small increase in ciliary length, which saturates as speed is increased further. We further verified detailed model predictions in the present case of a reduced motor speed. First, the model predicts that a slower IFT process should result in a lower IFT injection rate (Figure 4B), which is consistent with our data (Figure 3C). Second, in the case of the slower motors, the rate of change in IFT injection rate should decay faster than in wild-type (Figure 4C). To test this, we analyzed the ciliary length-dependent reduction in IFT injection rate in mutant and control cells during ciliary elongation. Indeed, the mutant cells exhibited a faster rate of reduction in IFT injection rate during earlier stages of ciliary assembly relative to the control (Figure 4D-E). We hypothesize that the faster reduction in IFT injection rate might be caused by the higher level of motor/IFT proteins resulted from the slower motor during ciliary assembly. To test this hypothesis, we measured the levels of ciliary IFT proteins in shorter growing cilia and found that the mutant cilia indeed had higher ciliary levels of IFT proteins (Figure 4F). Third, the ciliary regeneration kinetics predicted by the model for the two motor speeds are also consistent with our data (Figure 4G and H). Therefore, we have presented a simple physical model which reproduces the mild change in ciliary length upon a significant reduction in IFT velocity. This nonlinear dependence seems to be resulted from a combination of a limited pool of IFT resources and a separation in diffusive versus ballistic transport timescales for the IFT motor.

**Figure 4.**
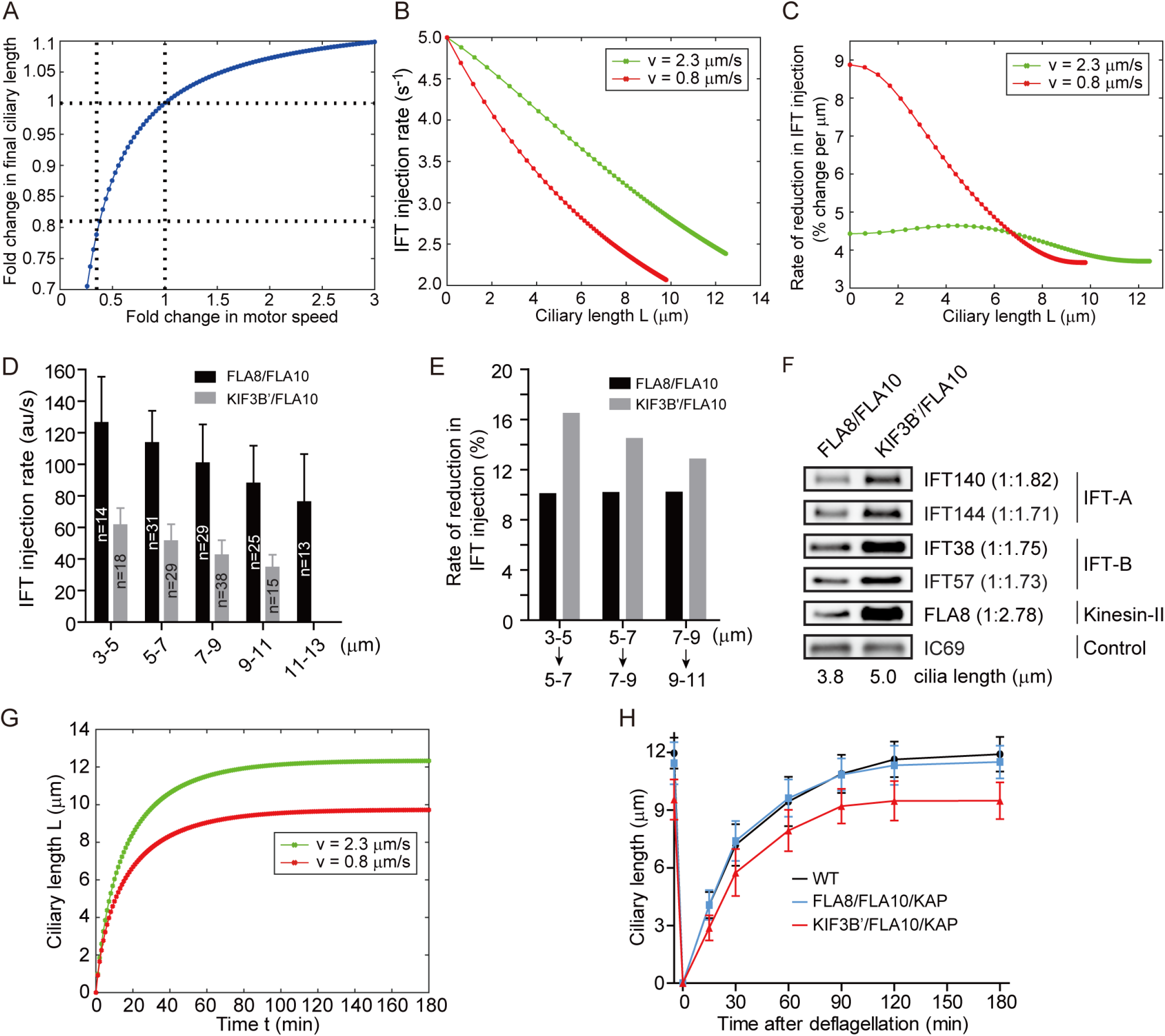
Mathematical modeling predicts a non-linear scaling relationship between motor velocity and ciliary length (See also Figure S5, 6). (A-C) Modeling simulation. Relationship between motor speed and steady state ciliary length (A); between ciliary length during ciliary assembly and IFT injection rate (B), and rate of reduction in IFT injection rate as a function of ciliary length (C). See Methods for detail. (D) IFT injection rate during ciliary assembly at different length of cilia as indicated. The fluorescence intensity of IFT46-YFP was monitored via TIRF microscopy. (E) Reduction in IFT injection rate along with ciliary elongation during ciliary assembly. The data were derived from (D). (F) Cells expressing slower chimeric motor KIF3B’/FLA10/KAP at shorter growing cilia exhibit higher ciliary levels of IFT proteins relative to the control. Isolated cilia during ciliary regeneration from cells as indicated were subjected to immunoblotting with the indicated antibodies. (G) Simulated kinetics of ciliary assembly during ciliary regeneration. (H) Kinetics of ciliary assembly. Cells were deflagellated by pH shock to allow cilia regeneration. Cells were fixed at the indicated times followed by measurement of ciliary length. Data shown are mean ± SD (n = 50).

## DISCUSSION

### IFT motor speed and ciliary length control

#### Motor limitation is a major determinant of ciliary length

Uniquely, our experimental system allowed us to evaluate how a single parameter change in motor speed affects ciliary assembly and length. Physical models have been proposed to explain ciliary length control (Fai et al., 2019; Hendel et al., 2018). However, they did not evaluate how motor speed influences ciliary length. We adapted the single-cilium model of Fai and colleagues with extended analysis to understand how motor speed affects length (Fai et al., 2019). These analyses suggest that motor speed should not significantly affect ciliary length even with a motor with ~2.8 fold reduction (Fig. 4G), which is consistent with our data (Fig. 4H). Based on this, we suggest that motor limitation is a key determinant of ciliary length control.

How could this constraint arise? The conventional view suggests kinesin-II diffuses back to the ciliary base during retrograde IFT (Chien et al., 2017; Engel et al., 2009; Mueller et al., 2005; Pedersen et al., 2006); and this diffusion was proposed to delay the return of kinesin-II motors, which would deplete the amount of kinesin-II available for anterograde transport (Chien et al., 2017). In this diffusion-limited scenario, a greater number of motors would lead to a greater IFT injection rate (Eq. 1) and, in turn, a longer steady-state ciliary length when the diffusion rate is constant. Thus, the diffusion timescale has been suggested to serve as a proxy by which cells are able to measure the length of their cilia (Hendel et al., 2018). The existence of a limited supply of motors would constrain the amount of IFT-associated proteins (e.g. tubulin) entering into cilia, which naturally entails a decreasing ciliary assembly rate when cilium is growing and confers a non-linear scaling between the IFT injection rate and the ciliary length (Eq. 3). Finally, the ciliary assembly rate is balanced with the disassembly rate, leading to a steady state ciliary length (Engel et al., 2009; Liang et al., 2014; Marshall et al., 2005; Marshall and Rosenbaum, 2001; Wemmer et al., 2020).

In this study, we showed that when IFT motors are slow, IFT injection rate is not only limited by diffusion but also by motor speed. As shown in (Figure S5A, B), when the motors are slow and the diffusion rate is unchanged, the number of motors needed to grow a cilium for a wild-type length is greater. In other words, when the number of motor and diffusion rate are both unchanged, slow motors would always lead to a shorter ciliary length (Figure S5B). Therefore, the reduction in motor speed, as we see in the mutant with a chimeric slow motor, switches the ciliary length control system from a diffusion-limited regime to a motor-limited one, which would lead to a reduction in IFT injection rate. In this scenario, the non-linear scaling relationship between the IFT injection rate and the ciliary length, which is intrinsic in our model, accounts for the mild reduction in ciliary length despite a 2.8-fold reduction in IFT velocity (Fig. 4A).

### Model implications and limitations

In the chimeric motor mutant, there should be more motors moving along the cilium (i.e. a greater *N*_*ballistic*_) because motors are slow, but there will be fewer motors diffusing back (i.e. a smaller *N*_*diffusive*_) because the cilium is shorter (Figure S6). These two changes have opposite effects on the total number of motors available for ciliary assembly and they cancel each other to a large extent. The net effect is that the number of motors available at the ciliary base in the mutants with slow motors becomes smaller, leading to a reduction in IFT injection rate. The lower IFT injection rate compensates for the longer ballistic time of the slower motors. The difference in *N*_*ballistic*_ + *N*_*diffusive*_ saturates, which explains the observation of the nearly unchanged level of motors in the cilium at the final steady state (Figure 3G). However, the difference is more pronounced in earlier stages of ciliary assembly when the cilia are short, so the ciliary length control is neither diffusion-limited nor motor-limited. In this scenario, slow motors are expected to result in a higher level of IFT motors and proteins in cilia (Figure S6). This is fully consistent with our observations (Figure 4F).

On the other hand, although our model recapitulates the net reduction in IFT injection rate in the slow-motor mutants, the expected level of change in our modeling results is smaller than that observed in our experiments. We think there are probably two reasons. First, the fold-change as observed in the experiments may be over-estimated because we may only have measured the upper bound of the fold change in IFT injection rate due to the limitations of fluorescence microscopy. Second, there could be a negative feedback mechanism that auto-regulates IFT injection frequency based on the amount of IFT complexes inside the cilium, which was not considered in our model. What could be the negative feedback mechanism? We speculate that the possible candidate mechanisms may involve signaling events at the ciliary base. For example, it has been shown that IFT injection is influenced by signaling such as Ran activation or FLA8/KIF3B phosphorylation (Liang et al., 2018; Ludington et al., 2013). Thus, abnormal higher level of motor/IFT complexes during ciliary assembly may invoke such signaling to restrict IFT entry, which remains to be a hypothesis awaiting for future studies. In addition, it has been reported that the occupancy of the ciliary cargoes on IFT complexes, which contributes to ciliary assembly and length, is regulated (Craft et al., 2015; Pan and Snell, 2014; Wren et al., 2013), which was not considered in our model. Our finding that a higher than expected reduction in IFT injection rate in the slower motor mutants did not significantly change ciliary length may also imply that the slower motor carries relatively more ciliary cargoes, e.g. tubulins, in each IFT train. This is an interesting point to be further tested.

#### Further remarks

Several mutants of *Chlamydomonas* with abnormally long cilia (up to 2-fold) have been identified (Asleson and Lefebvre, 1998). The underlying mechanisms remain elusive (Wemmer et al., 2020). Our model (Figure 4A) suggests that cilia length saturates with increasing motor speed, therefore the long cilia phenotype cannot be due to a change in anterograde IFT velocity. Indeed, it has been shown that the velocity of anterograde IFT is similar between long cilia mutant *lf4* and wild type cells (Wang et al., 2019). Interestingly, it was shown that a cilium elongation phenotype induced by the activation of PKA or depletion of ICK in mammalian cells is accompanied with an increased velocity of anterograde IFT (Besschetnova et al., 2010; Broekhuis et al., 2014). However, this increase in anterograde IFT velocity was thought to be a consequence, but not the reason, of cilium elongation, as IFT velocity itself was reported to be dependent on ciliary length (Engel et al., 2009).

### Kinesin-II and the ciliary entry of IFT complexes

The conventional view for IFT is that kinesin-II carries IFT complexes with their associated cargoes into cilia from the ciliary base through the transition zone (Prevo et al., 2015; Rosenbaum and Witman, 2002; Scholey, 2013; Wingfield et al., 2017). However, this view was challenged by arguments that IFT trains are picked up by kinesin-II only after they enter cilia via kinesin-II independent mechanism (Nachury and Mick, 2019). We showed that reduction in motor activity of the chimeric Crkinesin-II with human motor domains resulted in a similar reduction in ciliary entry of IFT trains, suggesting that kinesin-II likely carries IFT trains to enter cilia. However, our data do not preclude the possibility that kinesin-II activity indirectly affects IFT entry. For example, the slower velocity of the motor may alter the flux of kinesin-II motors reaching the distal transition zone where IFT motors and complexes are proposed to be assembled (Yang et al., 2019).

### Distinct mechanisms underlying formation of a functional kinesin-II for IFT from *Chlamydomonas* to mammals and *C. elegans*

The heterotrimeric organization of kinesin-II is essential for its function in IFT from lower eukaryotes to human (Scholey, 2013). This function requires a functional motor with the ability to bind the IFT complexes. The non-motor subunit KAP is required for full activation and targeting of kinesin-II to the ciliary base while KIF3B/FLA8 is required for binding the IFT complexes (Funabashi et al., 2018; Mueller et al., 2005; Sonar et al., 2020). We have revealed that the mechanism involved in forming this essential complex is distinct in *Chlamydomonas* compared to mammals and *C. elegans*. CrKinesin-II with two identical motor domains could coordinate *in vitro*, which is the same as in mammals and *C. elegans* (Brunnbauer et al., 2010; Muthukrishnan et al., 2009; Pan et al., 2010). However, such a chimera functioned *in vivo* in *Chlamydomonas* but not in *C. elegans* (Pan et al., 2010). The two motor subunits in *C. elegans* cannot form dimers and the particular form of the heterodimer is essential for association with KAP (Brunnbauer et al., 2010). In mammalian cells, KIF3A can self-interact and is able to interact with KAP while KIF3B can interact with KAP but cannot form homodimer, thus formation of a heterotrimeric kinesisn-II is required for IFT (Funabashi et al., 2018). For CrKinesin-II, both FLA10 and FLA8 could self-interact. The FLA10 homodimer is able to interact with KAP, however, such a complex is expected not to be able to function in IFT because FLA8/KIF3B is required for binding IFT trains. In contrast, FLA8 forms homodimers but cannot interact with KAP, thus was unable to function in IFT as well because KAP is required for full activation of the motor and its targeting to the ciliary base. Thus, this work reveals the molecular basis underlying the necessity of CrKinesin-II to function in IFT, and also highlights distinct mechanisms from both mammals and *C. elegans*.

In summary, we show that although the requirement of the heterotrimeric organization of kinesin-II required for IFT is conserved, the underlying mechanisms forming this heterotrimeric complex are distinct among various organisms. Furthermore, our studies with chimeric kinesin-II that has reduced motor speed yield new insights into how motor speed regulates IFT and ciliary length. Our data suggest that controlling IFT entry and hence cargo loading appear to be the key determinant of ciliary length control.

## MATERIALS AND METHODS

### Strains and cell cultures

The wild type strain 21gr (mt+, CC-1690) was available from the *Chlamydomonas* Resource Center (University of Minnesota, St. Paul, MN). A *fla8* mutant was generated previously in this laboratory (Liang et al., 2014). For the transgenic strains used in this study, please refer to Table S1. Cells were cultured on 1.5% agar plates or in liquid M medium (Sager and Granick, 1954) at 23°C with aeration under a 14:10 hour light-dark cycle.

### DNA constructs of chimeric kinesins for *in vitro* motility assay

Full-length cDNAs of KIF3A and KIFAP3 were gifts of Dr. Jiahuai Han (Xiamen University, China). Full-length cDNA of KIF3B was synthesized (WuXi Qinglan Biotech). FLA10, FLA8 and KAP cDNAs were cloned from a *Chlamydomonas* cDNA library (Takara). For chimeric kinesins, the motor domain of FLA10 was replaced with that of FLA8 or KIF3A to generate FLA8’ or KIF3A’ as specified in the text, respectively. The constructs for chimeric kinesins FLA10’ and KIF3B’ were similarly generated. The cDNAs with tags as indicated in the text were cloned in the pOCC vectors, respectively, by conventional molecular techniques.

### Protein expression and purification

Proteins used for *in vitro* studies were expressed in insect Sf9 cells using the baculovirus expression system. MBP-tag or His-tag at the C-terminus of the indicated proteins was used to facilitate purification while RFP-tag or GFP-tag at the C-terminus was used for imaging. The infected cells were grown for 3 days at 27 °C. Cells from 500 ml of cultures were disrupted by mortar and pestle grinding on ice in 100 ml lysis buffer (80 mM Pipes, pH 6.9; 150 mM KCl, 1 mM MgCl_2_, 1 mM EGTA, 0.1 mM ATP, 0.1% Tween-20). The cell lysates were centrifuged at 444,000xg for 40 min and 4 °C. HsKinesin-II was purified using Ni column and MBP column successively (Ni-NTA agarose affinity resin, QIAGEN; Amylose resin, NEB New England Biolabs). For purification of Crkinesin-II, the heterodimer purified from a MBP column and KAP from a Ni column were mixed, followed by purification via Superose 6 (GE Healthcare). The proteins were frozen in liquid nitrogen and stored at −80 °C.

### *In vitro*, single-molecule motility assay

A previously published protocol was followed for the *in vitro* motility assay (Gell et al., 2010). Briefly, 6.25 μl of 40 μM porcine brain tubulin mix containing 5% Alexa 647-labeled tubulin in BRB80 buffer with addition of 4% DMSO, 4 mM MgCl2 and 1 mM GTP (final concentrations) was incubated on ice for 5 min. Tubulins were allowed to polymerize for 2 hours at 37 °C. The reaction was stopped by adding 200 μl of warm BRB80 buffer containing 20 μM taxol. Microtubules were collected in the taxol-BRB80 buffer after Airfuge centrifugation. For motility assay, the taxol stabilized microtubules were attached to a cover glass surface coated with anti-tubulin antibodies followed by the addition of indicated purified kinesins. The samples were imaged by TIRF microscopy (Olympus IX83 equipped with an Andor 897 Ultra EMCCD). The data were processed by imageJ.

### Pull-down assay

To determine possible homodimer formation of FLA10 or FLA8, FLA10-MBP/FLA10-GFP or FLA8-MBP/FLA8-GFP that were cloned respectively in pEGFP-C3 vectors were co-expressed in HEK293T cells with controls as indicated in the text. The transfected cells after growing for 48 h were lysed in 500 μl lysis buffer (PBS, pH 7.4, 150 mM KCl, 1 mM MgCl_2_, 1 mM EGTA, 0.1 mM ATP, 0.5% NP-40) containing protease inhibitor cocktail. After 30 min on ice, the cell lysates were centrifuged at 20, 000xg for 10 min. The supernatant was mixed with anti-GFP beads and incubated at 4 °C for 2 h with constant rotation followed by washing with lysis buffer for three times. The samples were finally analyzed by immunoblotting with the indicated antibodies. To examine possible interactions of KAP with FLA10 or FLA8, FLA8-RFP/KAP-GFP-His or FLA10-MBP/KAP-GFP-His were co-expressed in Sf9 cells, respectively, with FLA8-RFP/FLA10-MBP/KAP-GFP-His as control. The transfected cells were lysed in lysis buffer (80 mM Pipes, pH 6.9; 150 mM KCl, 1 mM MgCl_2_, 1 mM EGTA, 0.1 mM ATP, 0.1% Tween-20, 10 mM imidazole) containing protease inhibitor cocktail. The proteins were pulled down by Ni beads followed by washing and immunoblotting with the indicated antibodies.

### Ectopic gene expression in *Chlamydomonas*

*FLA8-HA or FLA8-YFP was* cloned in between PSAD promoter and terminator in a modified vector pKH-IFT46 (kindly provided by Dr. Kaiyao Huang, Institute of Hydrobiology) that harbors hygromycin B resistance gene. The final construct was linearized with ScaI and transformed into the *fla8* mutant by electroporation (Liang and Pan, 2013). The construct of KIF3B’ for expression in *fla8* was made by replacing the motor domain of FLA8 with that of KIF3B. IFT46-YFP was provided by Dr. Kaiyao Huang (Lv et al., 2017).

### Ciliogenesis and ciliary assays

Cilia isolation or ciliary regeneration was performed as described previously (Wang et al., 2019; Zhu et al., 2017b). For ciliary regeneration, cells were deflagellated by pH shock to allow ciliary regeneration at the indicated times followed by fixation with 1% glutaraldehyde. Cells were imaged by differential interference contrast microscopy with a 40x objective on a Zeiss Axio Observer Z1 microscope (Carl Zeiss) equipped with an EM CCD camera (QuantEM512SC, Photometrics). Ciliary length from 50 cells at the indicated times was measured using ImageJ (NIH). For cilia isolation, control cells or cells during ciliary regeneration were deflagellated by pH shock. Sucrose gradient centrifugation was used to further purification of the detached cilia. Purified cilia were suspended in HMDEK buffer (50 mM HEPES, pH 7.2; 5 mM MgCl_2_, 1 mM DTT, 0.5 mM EDTA, 25 mM KCl) containing EDTA-free protease inhibitor cocktail (mini-complete, Roche), 20 μM MG132 and 25 μg/ml ALLN, frozen in liquid nitrogen and finally stored at −80 °C until use.

### SDS-PAGE and immunoblotting

Analysis for SDS-PAGE and immunoblotting has been described previously (Wu et al., 2018). Cells were collected by centrifugation and lysed in buffer A (50 mM Tris-HCl, pH 7.5; 10 mM MgCl_2_, 1 mM DTT and 1 mM EDTA) containing EDTA-free protease inhibitor cocktail (mini-complete, Roche), 20 μM MG132 and 25 μg/ml ALLN followed by boiling in SDS sample buffer. Proteins separated on SDS-PAGE were analyzed by coomassie blue staining or immunoblotting.

Rabbit polyclonal antibodies against IFT57 and IFT38 were made by immunizing polypeptide 1-260 aa and 275-443aa, respectively, and affinity purified (Abclone, China). The other primary antibodies were detailed in Table S2. The HRP-conjugated secondary antibodies were the following: goat anti-rat, goat anti-rabbit and goat anti-mouse (1:5000, EASYBIO, China).

### Live cell imaging of IFT

Total internal reflection fluorescence (TIRF) microscopy was used to observe live IFT. The coverslips treated with 0.01% (v/v) polylysine (Sigma) were used to immobilize cells. Images were acquired at room temperature on a Nikon microscope (A1RSi) equipped with a 100x (N.A. 1.49) TIRF objective and a cooled electron-multiplying CCD camera (Orca-flash 4.0; Hamamatsu, Japan). Images were analyzed with ImageJ (NIH, USA). The IFT speed, IFT frequency and IFT injection rate were measured following previous publications (Engel et al., 2009; Wemmer et al., 2020). The number of anterograde fluorescent IFT trains entering cilium per unit time was calculated for IFT frequency. To obtain the IFT injection rate, the fluorescence intensity of IFT trains (normalized for camera noise) per unit length of cilium and the velocity of anterograde IFT were first measured. Because most IFT trains do not stop during the transport, IFT injection rate was then calculated as the product of the fluorescent intensity and the velocity.

### Thin-section electron microscopy

Previously published protocols were followed (Craige et al., 2010; Meng et al., 2014). The samples were imaged on an electron microscope (H-7650B; Hitachi Limited) equipped with a digital camera (ATM Company).

### Mathematical modeling

We adapt a recently published model (Fai et al., 2019; Hendel et al., 2018) to understand the mild-reduction in cilium length upon 3-fold reduction in IFT velocity. Here, kinesin-II motors transport anterograde IFT trains with speed *v*, deposit cargoes including tubulin subunits at the growing cilium tip and then diffuse back to the base (with diffusion constant *D*). As proposed in Fai and colleagues (Fai et al., 2019), we assume that over cilia regeneration timescales, there is a limiting total pool of motors (*N*) and of tubulin (*T*) per cilium. Without taking into account the detailed mechanism of the IFT injection, the IFT flux or IFT injection rate (they are the same given that IFT trains rarely stop during transport) (*J*) is considered to be proportional to the number of free motors *N*_free_ (motors that are neither moving along the cilium nor diffusing back to the base) with a kinetic constant *K* (Eq. 1).

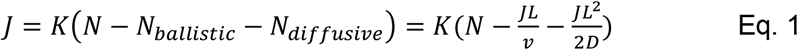

Here, we have approximated the dynamics as a quasi-steady state process in which the ballistic and diffusive fluxes are balanced – with the ciliary tip acting as a diffusive source and the base as a sink. This is because the timescale for transport of IFT particles over the length of the cilium (seconds) is much less than that of cilium regeneration (hours). Rearranging Eq. 1, we obtain

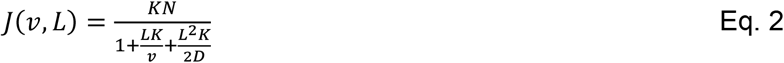

which reproduces the empirical finding that *J* decreases with increasing cilium length (Engel et al., 2009). Moreover, the formula predicts that *J* should decrease with decreasing v, when all other parameters are held constant. Both of these features are consistent with our findings (Figure 4B). Note that the scaling in Eq. 2 applies to any rate-limiting IFT protein (not only motors). Furthermore, the net assembly rate of cilium can be given by

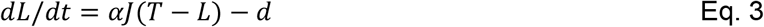

where the additional constant α is a tubulin binding factor, and *d* is a constant disassembly rate. This reproduces the empirical observation that the net assembly rate is decreasing as the cilium grows longer (Marshall et al., 2005; Marshall and Rosenbaum, 2001).

Finally, combining Eq. 2 and Eq. 3, we could obtain one positive solution (provided *T>d/αKN*) for the final steady-state ciliary length (Eq. 4)

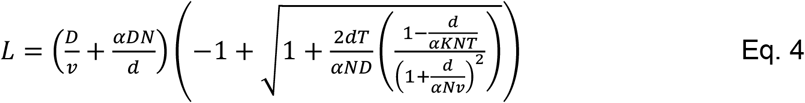

Using Eq. 2 and Eq. 4, we evaluated the scaling relationship between the IFT velocity and ciliary length using numerical simulations with the following choice of parameters (*α* = 10^−4^, *K* = 5 min^−1^, *N* = 60, *D* = 10 *μm*^2^/*s*, *T* = 30 *μm*, and *d* = 0.25 *μm*/*min*). These parameters are representative of measured values obtained from the literature (Fai et al., 2019; Hendel et al., 2018) and have been chosen to produce a final length of approx. 12 μm at the wild type motor speed. Variations within a realistic range of these values were found to have little effect on the overall functional dependencies, usually only resulting in faster or slower growth kinetics and/or a different final cilium length.

We further explored the dependence of ciliary growth on physiological model parameters for motors with normal and reduced speed (Figures S5). At normal motor speeds, the final length of cilium is diffusion-limited. In contrast, a slow motor will always lead to a shorter than wild-type cilium regardless of the rate of diffusion unless the number of motors available in circulation is increased (motor-limited). Thus, a motor with slower speed would limit IFT entry due to motor limitation (Figure S5A, B). The situation for tubulin limitation is similar (Figure S5C, D). Meanwhile, we found little difference between the cilia growth timescales in the case of the wild-type motor speed, compared to the ~3x slower motor (Figure S5E, F). This is consistent with the data (Figure 4H).

### Quantification and statistical analysis

Independent experiments were carried out for at least two or more times. Data plotting was performed using Prism (GraphPad7). The data were presented as mean ± SD. Statistical significance was performed by using two-tailed Student’s t test analysis. p< 0.05 was considered to be statistically significant. *, p<0.05; **, p<0.01; ***, p<0.001; ****, p<0.0001.

## ACKNOWLEDGMENTS

We are grateful to Dr. Jonathon Howard (Yale University) for discussion during the course of this work. This work was supported by the National Key R&D Program of China (2018YFA0902500, 2017YFA0503500) and the National Natural Science Foundation of China (31991191, 31671387, 31972888) (to J.P.).

## AUTHOR CONTRIBUTIONS

S.L., W.C., H.T., X.L., and J.P. designed the experiments and analyzed the data; S.L., W.C. and H.T. performed research; K.W. analyzed the data and performed modeling analysis; J.P., X.L., and K.W. contributed reagents and analytic tools; J.P., X.L., K.W. and S.L. wrote the paper.

## DECLARATION OF INTERESTS

The authors declare no conflict of interest.

## SUPPLEMENTAL INFORMATION

Supplemental information include two tables and six figures.

## SUPPLEMENTAL INFORMATION

**Figure S1.**
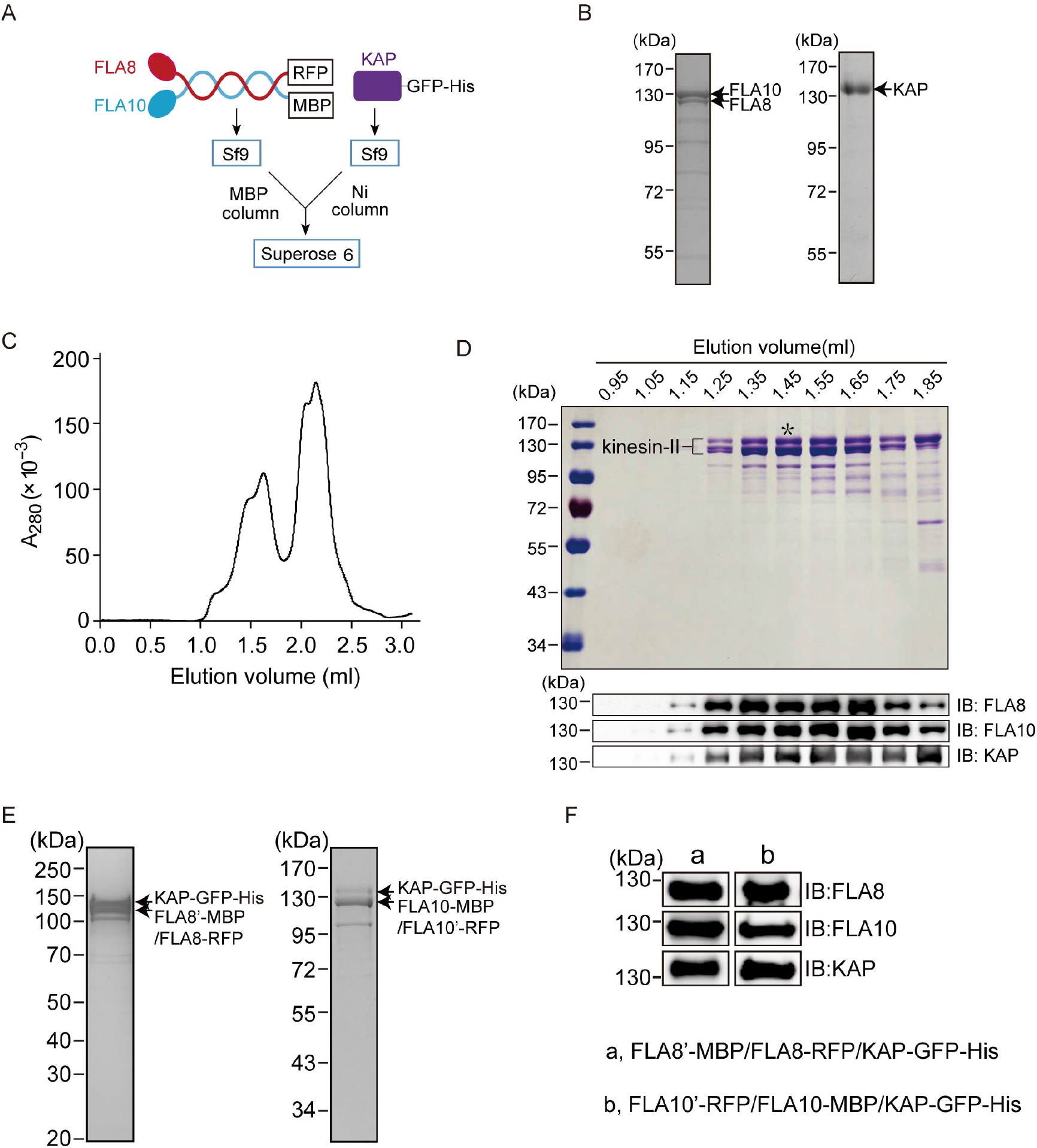
Purification of recombinant CrKinesin-II and chimeric Crkinesin-II with identical motor domains. Related to Figure 1. (A) Strategy for expression and purification CrKinesin-II. (B) Coomassie-stained SDS-PAGE of purified FLA10/FLA8 from an MBP column (left) and KAP from a Ni column (right). (C) Elution profiles of CrKinesin-II from gel filtration. Please refer to Figure S1D. (D) Analysis of purified CrKinesin-II by SDS-PAGE and immunoblotting. Elution fractions of CrKinesin-II from gel filtration were separated on a 10% SDS-PAGE followed by coomassie blue staining (top) or by immunoblotting with the indicated antibodies (bottom). An asterisk indicates the fraction used for motility assay. (E-F) FLA8’-MBP/FLA8-RFP/KAP-GFP-His and FLA10’-RFP/FLA10-MBP/KAP-GFP-His were expressed respectively in Sf9 cells followed by purification using MBP and Ni affinity columns. The purified products were analyzed by SDS-PAGE followed by coomassie blue staining (E) or immunoblotting (F).

**Figure S2.**
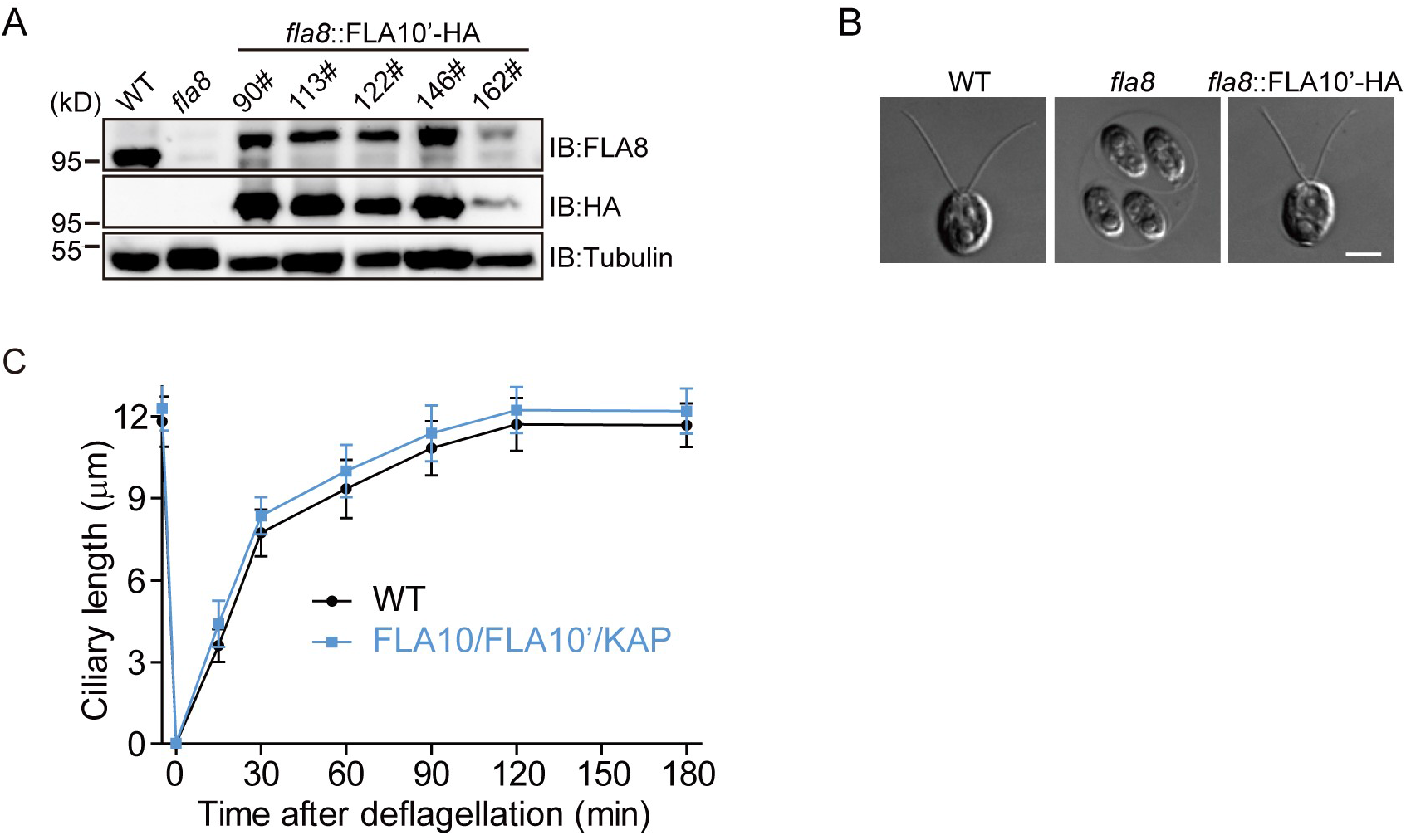
Cells expressing chimeric CrKinesin-II with identical motor domains of FLA10 form normal cilia. Related to Figure 1. (A) HA-tagged FLA10’ (motor domain of FLA10 with FLA8 carboxyl tail) was transformed into an aflagellate mutant fla8. The transformants were expected to form a chimeric motor with two identical motor domains of FLA10. The transformants were examined by immunoblotting with wild type (WT) and fla8 cells as control. (B) FLA10’ rescued the aflagellate phenotype of fla8. Shown are the differential interference contrast images of the cells as indicated. Bar, 5 μm. (C) Ciliary regeneration kinetics of FLA10’ transgenic strain with wild type cells as a control. The cells were deflagellated by pH shock to allow ciliogenesis. Ciliary length of the cells was measured before deflagellation or at the indicated times after deflagellation. Data shown are mean ± SD (n = 50).

**Figure S3.**
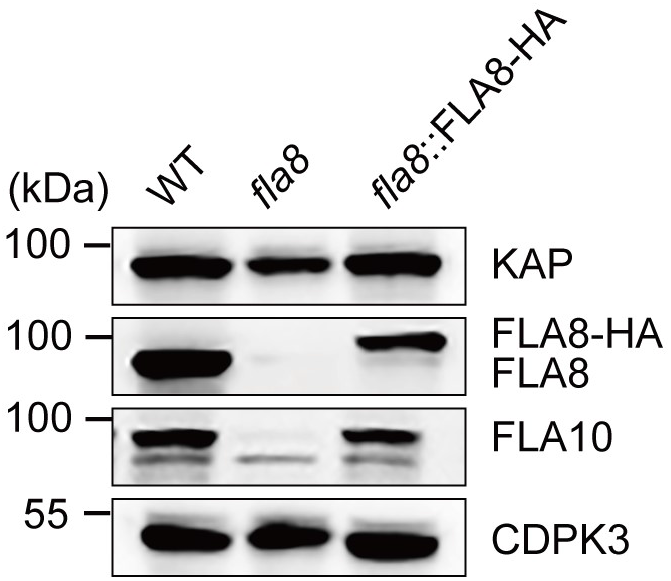
Loss of FLA8 in a fla8 mutant results in diminished level of FLA10. Related to Figure 1. Wild type (WT), fla8, and rescued strain of fla8 were analyzed by immunoblotting with antibodies against CrKinesin-II subunits KAP, FLA8, FLA10, and CDPK3 (Calcium dependent kinase 3) as a control.

**Figure S4.**
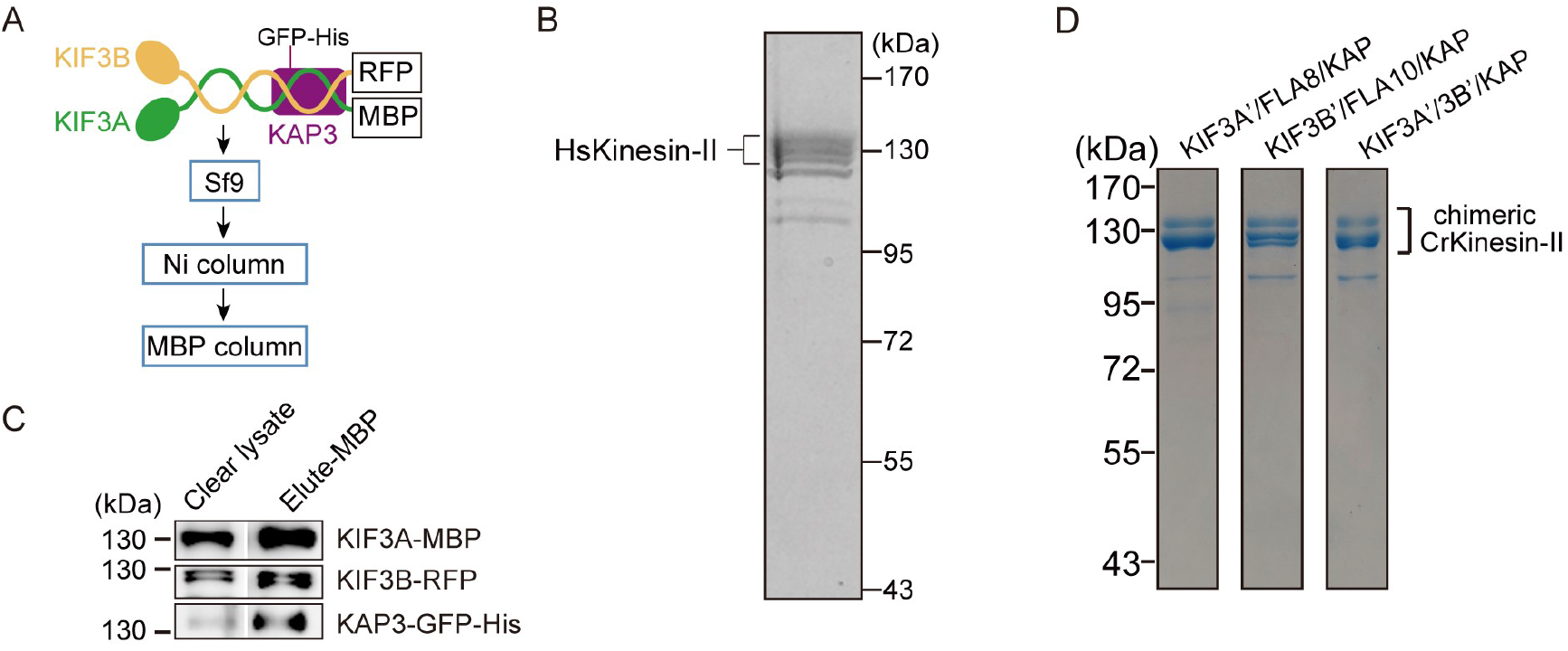
Purification of recombinant HsKinesin-II and chimeric CrKinesin-II with motor domains of HsKinesin-II. Related to Figure 2. (A-C) Expression and purification of recombinant HsKinesin-II. Strategy of expression and purification (A). Purified kinesin-II were subjected to SDS-PAGE analysis followed by coomassie blue staining (B) or immunoblotting with antibodies against KIF3A, KIF3B or KAP3 (C). (D) Purification of chimeric CrKinesin-II with motor domains of HsKinesin-II. The motor domains of recombinant CrKinesin-II as shown in Figure 1A was replaced with one or two motor domains of HsKinesin-II as indicated. The resulting constructs KIF3A’-MBP/FLA8-RFP/KAP-GFP-His, KIF3B’-RFP/FLA10-MBP/KAP-GFP-His and KIF3A’-MBP/KIF3B’-RFP/KAP-GFP-His were expressed respectively in Sf9 cells followed by purification using MBP and Ni affinity columns consecutively. The purified products were analyzed by SDS-PAGE followed by commassie blue staining.

**Figure S5.**
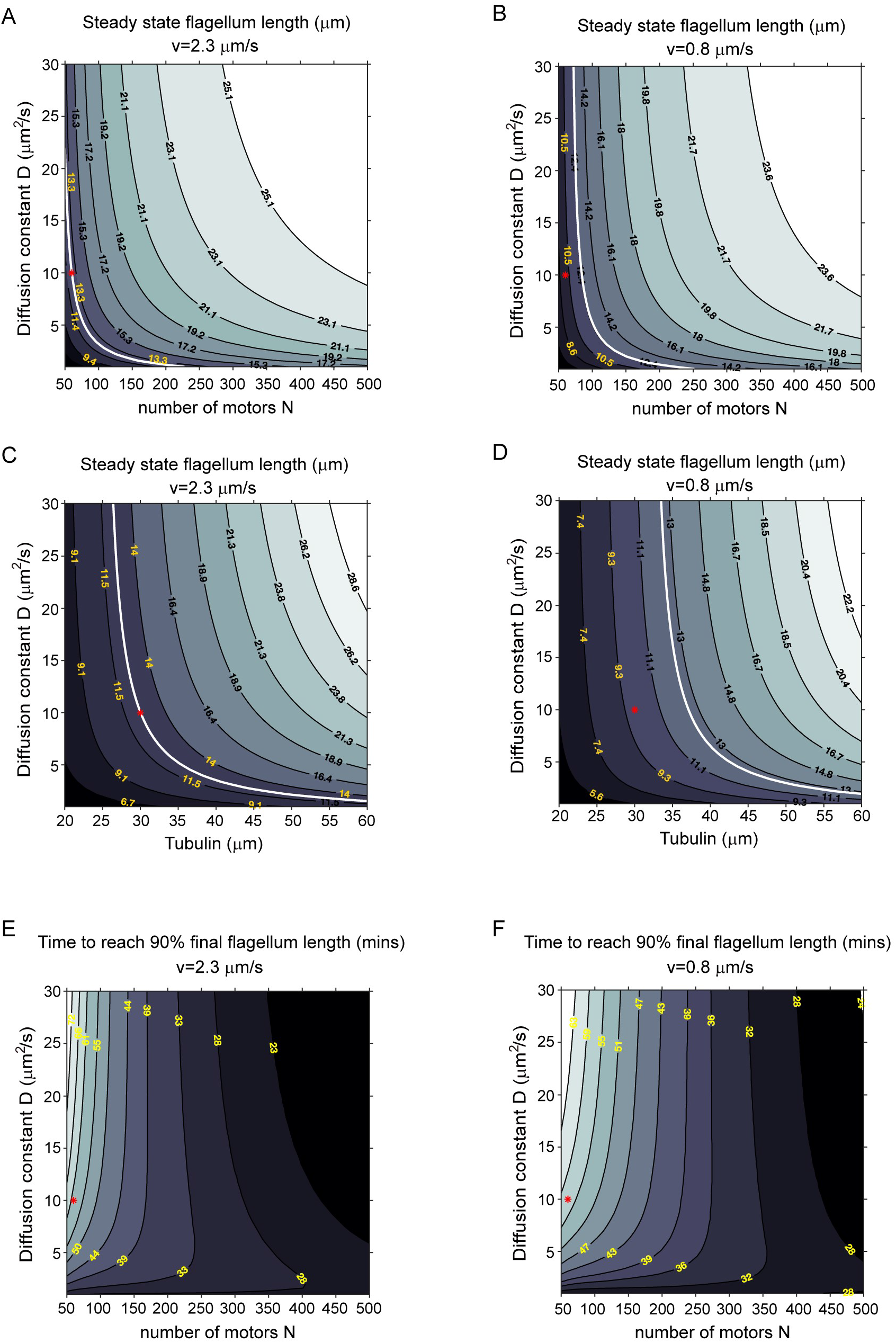
The number of motors and the amount of tubulins on ciliary length and the timescale to reach final ciliary length predicted by the modelling. Related to Figure 4. (A-D) Contour plots of the steady state cilium length when either N, the total number of motors or T, the total amount of tubulin, is varied with the rate of diffusion (D), while keeping all other parameters constant. The white lines correspond to 12.5 μm – assumed to be the wild-type cilium length. Red markers denote parameters used in the simulations presented in the main text. (E, F) Contour plots of the growth time (in minutes) required to attain 90% of the final steady state flagellum length.

**Figure S6.**
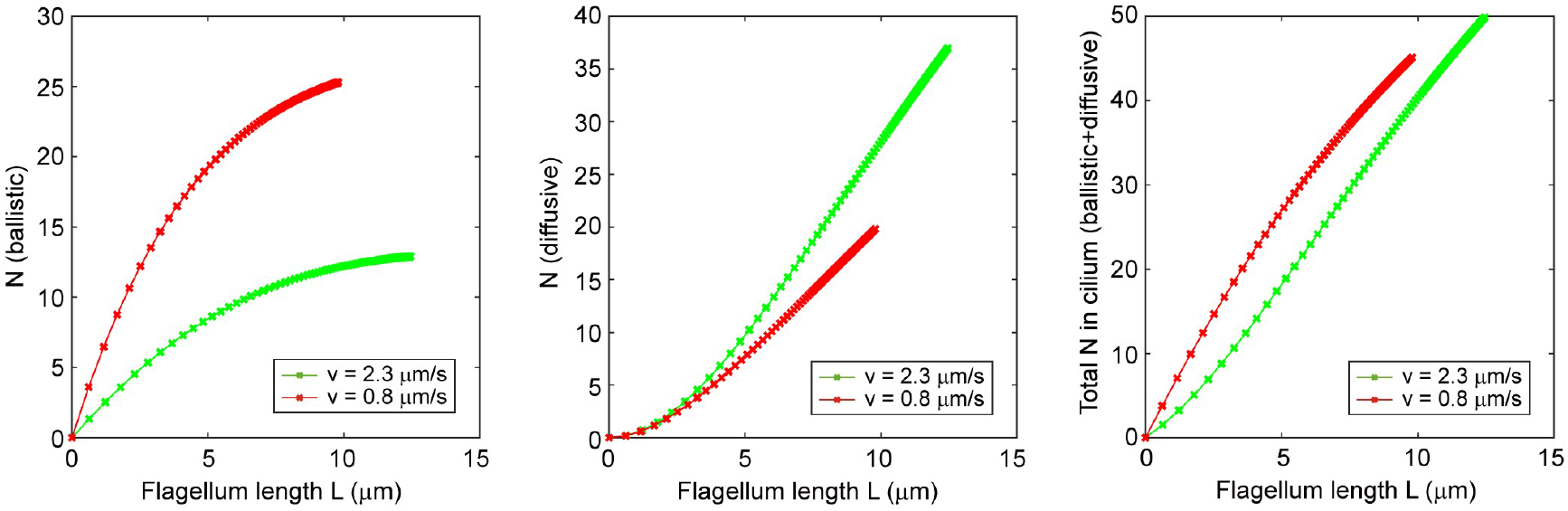
Model predictions of the number of motors inside the growing cilium. Related to Figure 4. Change in the number of motors found at each stage of the IFT cycle (ballistic, diffusive, combined) as the cilium grows. Representative plots are for parameters used in the simulations presented in the main text.

**Table S1.**
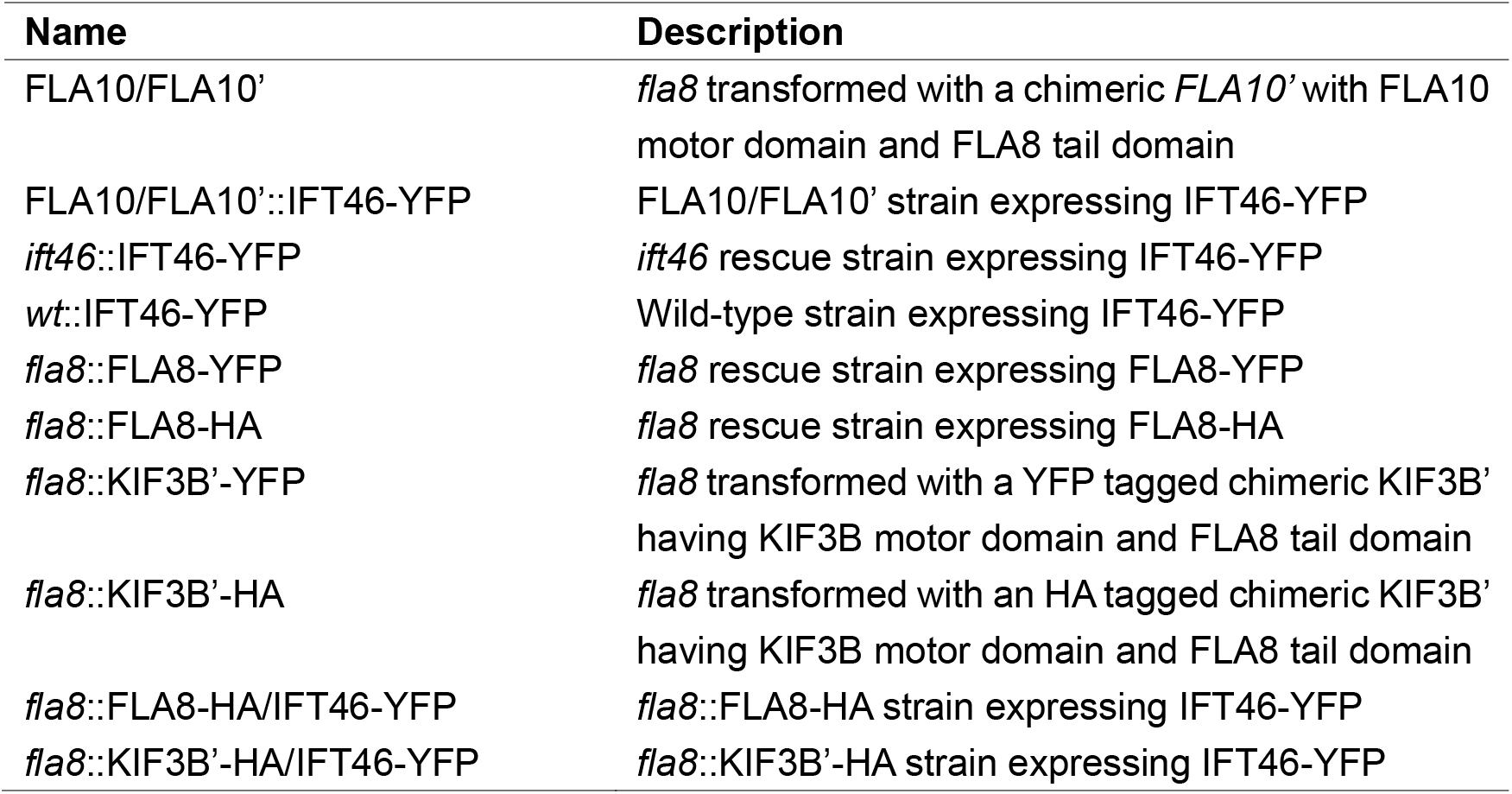
Transgenic strains used in this study.

**Table S2.** Primary antibodies used in this study.

| **Antibody** | **Dilution (IB)** | **Reference or source** |
| --- | --- | --- |
| Rat anti-HA | 1:1000 | Roche |
| Mouse anti-MBP | 1:1000 | CMCTAG |
| Mouse anti-GFP | 1:2000 | Abmart |
| Mouse anti-α-tubulin | 1:3000 | Sigma-Aldrich |
| Mouse anti-IC69 | 1:20000 | Sigma-Aldrich |
| Rabbit anti-IFT140 | 1:2000 | (Zhu et al., 2017) |
| Rabbit anti-IFT144 | 1:2000 | (Zhu et al., 2017) |
| Rabbit anti-IFT46 | 1:3000 | (Lv et al., 2017) |
| Rabbit anti-IFT38 | 1:3000 | This study |
| Rabbit anti-IFT57 | 1:2000 | This study |
| Rabbit anti-D1bLIC | 1:1000 | (Meng and Pan, 2016) |
| Rabbit anti-FLA8 | 1:3000 | (Liang et al., 2014) |
| Rabbit anti-FLA10 | 1:3000 | (Cole et al., 1998) |
| Rabbit anti-KAP | 1:3000 | (Liang et al., 2014) |
| Rabbit anti-KIF3A | 1:2000 | Abcam |
| Rabbit anti-KIF3B | 1:500 | Abcam |
| Rabbit anti-KIFAP3 (KAP3) | 1:5000 | Abcam |

